# P53 Transposable Elements and Regulatory Introns Inform Codondex Cell Selection for Autologous Trigger of Immune Cascade

**DOI:** 10.1101/508903

**Authors:** Kevin Bermeister

## Abstract

We were motivated to discover whether our small signatures triangulate patient cohort relationships between intron and mRNA or protein. We tested biological outcomes using the signatures and intend to use it to screen cells for use in autologous and allogeneic immunotherapies.

## Background

In and among genes, Transposable Elements (TE’s) include LINE, SINE, Retrotransposons and ALU repeats from the long evolutionary history of viral invasion, some of which remain very active^1^. Many TE’s co-opt roles in regulation of DNA including functions like unwinding, gene exposure and splicing site preference sometimes as a direct response to other cellular activities.

The dominance of TE’s throughout non-coding (NC) regions, predominantly among intragenic introns make TE classification important in understanding regulatory contribution to gene and chromosome function. DNA arrangements are more stable without TE’s because some have actively mobile tendencies. At least TE’s make DNA order a dynamic as influenced by epigenetic modifiers and factors that bind DNA response elements.

Initial chromosomal structure and function, after S and G phase can be considered causative since meiosis results in near perfect duplication at next generation cells. Therefore, new cells begin life pre-programmed with inherent NC-TE DNA encoded tendencies to respond to stimulus. Chromosomes are large molecules of DNA that can be described as a siloed concentration of fixed and mobile repetitive oligo arrangements.

Replicated, split or spliced a myriad independent DNA and RNA molecules are released from chromosomal bound DNA to represent free concentrations as part of a highly complex, sensitive dissociation. Innate equilibrium between chromosome and rest of cell may be governed by various mechanisms including histone driven exposure of DNA. These may present a form of response to global cellular stress where chromosomal bound and unbound DNA/RNA is inclined to an equilibrium by innate capacity to activate essential cell machinery.

Non-coding, intronic DNA sequences relative to coding mRNA or protein has been formulated as a specific view. Developed by Codondex it may describe aberrant encoding in regulatory mechanisms by comparing multiple patient data. Accordingly any change, even a single nucleotide represents a contribution to the upstream and corresponding potential cell dynamic.

Our big data analysis ranks DNA’s potential, for patient gene transcripts to be expressed into upstream representations including as RNA. Each intron DNA Element (DE) of more than 7 oligos is counted for its repetitive identical, same length DE’s. We hypothesize this can be used to represent the inherent siloed DE concentration signature of any given chromosome. Separately protein or mRNA is the actual measurement of an expression from the gene’s DNA coding regions known as exons.

Since many TE’s are predominantly intronic and mobile they can exist in different places in different patient DNA. In the analysis of patient transcripts, intron bound DNA and mRNA/protein signatures are considered to represent a relationship between potential and actual DNA-RNA states. Patient specific DNA as compounded by TE’s presents an enormous challenge for allogeneic therapy.

We developed a signature that compares a gene transcripts potential DNA (“pkmer”) anchored by its actual mRNA or protein to multiple patient cohorts. We intend using the signature to identify highly specific cell selections that will alert and activate Natural Killer cells sufficiently to trigger an immune cascade.

## Scientific Review

Codondex linked pkmer analysis through study of viral infection to TE’s, introns, their role in TP53, BRCA1, Natural Killer ligand selections and response. In its review of papers it found ample evidence in support - excerpts from this research follows:

**B**y the hypothesis, the specialized terminal structures of eukaryotic mRNA provide the ideal molecular environment for the lengthening, evolution, and subsequent massive expansion of highly mobile retrotransposons, leading directly to the retrotransposon-cluttered structure that typifies modern metazoan genomes and the eventual emergence of retroviruses^2^.

**H**owever, recent studies indicate that retention of specific intronic, non-protein-coding sequences within cytoplasmic mRNA (Cytoplasmic Intron sequence-Retaining Transcripts, CIRTs) in mammalian neurons and other cells plays a role in producing functional proteins^3^.

**H**ere, we show that downregulation of the mRNA of BRCA1 is associated with increased transcription of SINEs and production of sense and antisense SINE small RNAs. We find that BRCA1 mRNA is post-transcriptionally down-regulated in a Dicer and Drosha dependent manner and that expression of a SINE inverted repeat with sequence identity to a BRCA1 intron is sufficient for downregulation of BRCA1 mRNA^4^.

**O**ver-expression of wild-type p53 in cancer cell lines strongly up-regulated the expression of ULBP1 and ULBP2, but not of other NKG2DLs, upon binding of p53 to response elements located in the intronic regions of these genes^5^.

**E**BV miRNAs also target other cellular transcripts such as the natural killer (NK) ligand lectin-like transcript 1, importin 7, mitochondrial import receptor subunit TOM22 homolog, F-box only protein 9, caprin family member 2, the CAP-Gly domain-containing linker protein 1, mitochondrial GTPase GUF1, lysine-specific histone demethylase 4B, transcription regulator zinc finger protein 451 and deubiquitinating enzyme OTUD1. Subsequently, fourteen additional viral miRNAs are identified in EBV-positive effusion lymphoma cell lines, and all of these miRNAs are derived from a miRNA cluster located within introns of the BART gene^6^.

**n**c886 (=vtRNA2-1, pre-miR-886, or CBL3) is a newly identified non-coding RNA (ncRNA) that represses the activity of protein kinase R (PKR). nc886 is transcribed by RNA polymerase III (Pol III) and is intriguingly the first case of a Pol III gene whose expression is silenced by CpG DNA hypermethylation in several types of cancer. PKR is a sensor protein that recognizes evading viruses and induces apoptosis to eliminate infected cells. Like viral infection, nc886 silencing activates PKR and induces apoptosis. Thus, the significance of the nc886:PKR pathway in cancer is to sense and eliminate pre-malignant cells, which is analogous to PKR’s role in cellular innate immunity. Beyond this tumor sensing role, nc886 plays a putative tumor suppressor role as supported by experimental evidence. Collectively, nc886 provides a novel example how epigenetic silencing of a ncRNA contributes to tumorigenesis by controlling the activity of its protein ligand^7^.

**O**ur previous study demonstrated that DR4 expression could be regulated in a p53-dependent fashion. In the present study, we have demonstrated that DR4 is a p53 target gene and is regulated by p53 through a functional intronic p53 binding site (p53BS) based on the following lines of evidence: (*a*) the p53BS in the *DR4* gene is almost identical to the one found in the first intron of the *DR5* gene in terms of their locations and sequences; (*b*) DR4 p53BS bound to p53 protein in intact cells upon p53 activation as demonstrated by a chromatin immunoprecipitation assay; (*c*) a luciferase reporter vector carrying the DR4 p53BS upstream of an SV40 promoter exhibited enhanced luciferase activity when transiently cotransfected with a wild-type p53 expression vector in p53-null cell lines or stimulated with DNA-damaging agents in a cell line having wild-type p53; and (*d*) when the DR4 p53BS, together with its own corresponding promoter region in the same orientation as it sits in its natural genomic locus, was cloned into a basic luciferase vector without a promoter element, its transcriptional activity was strikingly increased by cotransfection of a wild-type p53 expression vector or treatment with DNA-damaging agents^8^.

**I**n contrast, CpG sites, located within the first intron, were predominantly methylated in sample H1, H7 and placenta^9^.

**B**y applying rigorous criteria, a list of genes with mutations that are implicated unequivocally in the pathogenesis of ALS can be generated. These genes can be grouped into several loose categories: genes that alter proteostasis and protein quality control; genes that perturb aspects of RNA stability, function and metabolism; and genes that disturb cytoskeletal dynamics in the motor neuron axon and distal terminal. The mutations involved are mostly missense substitutions, although the genetic lesion in C9orf72 is an enormous expansion of an intronic hexanucleotide repeat^10^.

**T**his novel regulatory element may have allowed the SINE to mediate transcription of the long isoform of Nkg2d directly from exon 1b, lifting the selection pressure to maintain a splice acceptor signal at the intron1–exon1b junction. The mutation of this splice acceptor site forces the transcript starting from exon 1a to splice directly to exon 2, thus creating NKG2D-S^11^.

**M**ore specifically, 96.3% of TEs enriched in gene bodies overlap introns, in line with the normally observed distribution of introns and exons in the human genome. Notably, the only tissues with an enrichment of young TEs (specifically primate specific) are blood related (Mononuclear and Lymphoblastoid Cells). These results are in agreement with an elegant study that discovered a key role of primate specific TEs in the regulatory evolution of immune response. TE families enriched in active regions across at least 20 of the 24 tissues correspond to DNA TEs and SINEs^12^.

**T**hus, lack of p53 function might enable cancer cells not only to promote their proliferation and survival but also to escape from recognition by NK cells and, possibly, other NKG2D-bearing effector cells such as CD8ķ T cells, NKT cells, and gdķ T cells by defective upregulation of certain NKG2D ligands. In addition, coculture of NK cells and wtp53-, but not mutp53-, expressing cancer cells resulted in the NKG2D-dependent production of IFN-g. Because NK cell–derived IFN-g is important for the induction of Th1 and Tc1 effector responses, lack of p53 function might also impair the NK cell–dependent priming of T cells. Thus, not only cancer cell intrinsic but also extrinsic, immune cell-mediated pathways are likely to be affected by the impairment of p53 function in tumor cells^13^.

**A**lthough NF-κB and IRF3, the two major factors for triggering the innate immune response, cooperate to control the expressions of a variety of cytokines and chemokines due to viral infection, and NF-_*κ*_B alone has been shown to modulate the p53-mediated apoptosis pathway, the activation of the NF-_*κ*_B might not be necessary for PML-dependent activation of p53 by IRF3. However, it is plausible that activation of both NF-_*κ*_B and IRF3 should accelerate PML-dependent p53 activation and tumor-suppressive function. In conclusion, our results indicate that direct transcriptional activation of PML by IRF3 results in the p53-dependent growth inhibition of cancer cells, which is suggestive of a novel regulatory network between the innate immune response and tumor suppression^14^.

**A**ll together, these studies suggest that p53 accumulation could represent a key determinant of the immunogenicity of stressed cells that are infected or undergoing malignant transformation through its ability to regulate innate immune activation^15^.

## Signature, Bio-Testing and Method

<u>In a previous paper</u> we posited that specific intron derived, TP53 RNA subsequence concentrations directly correlate p53 binding autonomous and non-autonomous response elements and suggest similar for BRCA1. In conjunction with Professor Noam Shomron Lab and Tel Aviv University we subsequently entered a series of experiments by transfecting HeLa Cells with specific short intron-1 sequence selections, identified by our algorithms to test our bioinformatic hypothesis.

Our multi-stage experimental regime has been designed to determine the utility of an autologous treatment protocol that will; screen patient intron-1 DNA and mRNA/protein from multiple samples of biopsied cells; select sample; co-culture selected cells with NK cells and; apply in a dosage controlled therapy.

We first considered whether relationships between intron-1 and protein were non-random. We imported publicly available transcript data from ensembl.org for each of multiple genes including TP53 and BRCA1. For each transcript we computed pkmers to represent every potential DE of every oligo length greater than 7. For each transcript pkmer we computed a proteo-genetic signature by associating it with a signature of the transcript protein. We established and ran a series of experiments comparing pkmer:protein signatures. We used two different methods to generate randomized pkmers for the transcript, each of which were associated with the constant transcript protein signature. Working with Professor Mark Kon, head of Statistics and Bioinformatics at Boston University and Professor Liran Carmel, head of Bioinformatics at Hebrew University, we confirmed these relationships were non-random.

We interrogated each pkmer to identify the individual computations that contributed most to non-randomness in all pkmer:protein signatures for the set of 15 TP53 and 29 BRCA1 transcripts. Our most ‘non-random’ bioinformatic selections identified intron-1 derived pkmer:protein sequences for the aggregate pkmers of all gene transcripts^16^.

We identified a short list of intron-1 pkmers for each of TP53 and BRCA1 which we considered to have contributed most significantly to the non-random character of their intron-1:protein signatures. The list represents the deduplicated set of selected pkmer sequences computed for each transcript. Based on a ranking methodology we selected, 8 x 28-oligo RNA sequences which were synthesized for transfection in HeLa.

## Methods

In the Shomron Lab at Tel Aviv University, Codondex RNA oligos entered cells via liposomal transfection. HeLa cells were seeded in 24-well or 6-well plates at a concentration of 8×104 cells/well or 3×105 cells/well respectively. The cells were transfected with 15 pmol for 24-well or 75pmol for 6-well of RNA, using Lipofectamine 2000 Transfection Reagent (Thermo Fisher Scientific, USA) according to the manufacturer’s instructions. Transfection efficiencies were measured by CY-3 fluorescence in all cells, indicating a transfection efficiency of a minimum of 50% repeatedly. RNA was purified from the cells 24-72h hours after transfection.

Codondex designed RNA oligos were elevated between 3000 and 700,000 fold in the cells. Sequence expression of all eight short RNA was upregulated, compared to scrambled sequence overexpression control.

Proliferation Fold Change (FC) following 72h of RNA overexpression (OE) in the HeLa cell-line. All short RNA OE affected the proliferation rate, and was significantly down-regulated in six out of the eight sequences compared to a scrambled sequence OE control. Paired Student’s t-test was used for the statistical analysis (n=3, *p<0.05, **p<0.005).

Expression of all six short RNA sequences was upregulated compared to scrambled sequence OE control. Codonex designed RNA oligos entered the cells and were elevated between 200 and 50,000 fold. Two RNA sequences were dropped given they did not show a significant cell proliferation affect above 200x elevation.

Reverse transcription reaction experiment for mRNA was conducted using the random-primer and High-Capacity cDNA Reverse Transcription Kit (Thermo Fisher Scientific, USA). Reverse transcription reaction for specific mature Small RNA was conducted using TaqMan Small RNA assays according to the manufacturer’s protocol (Thermo Fisher Scientific, USA). Single Small RNAs/mRNAs expression was tested similarly using the TaqMan Universal PCR Master Mix (Thermo Fisher Scientific, USA), or the SYBR green fast PCR master mix (Quantabio, USA), respectively.

## Results

The six Codonex RNA oligos modified TP53 and BRCA1 mRNA levels.

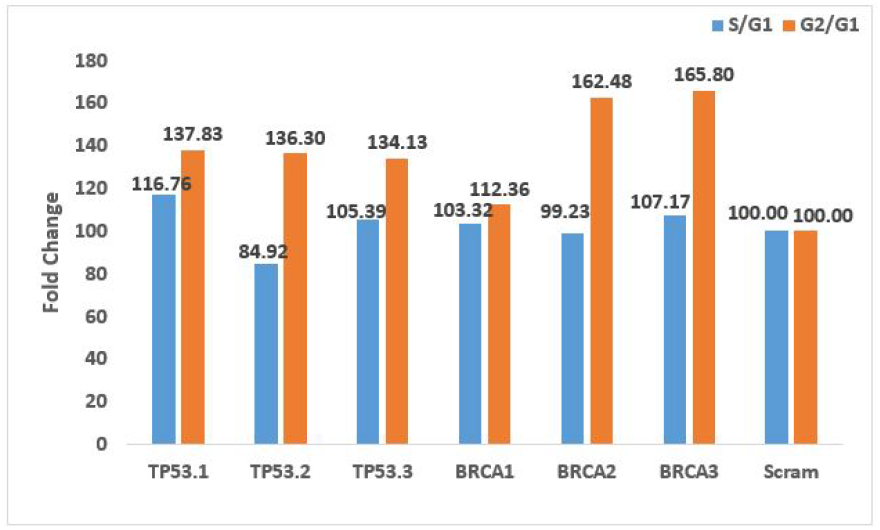

Cell-cycle analysis following 24h of RNA OE in the HeLa cell line. S/G1 or G2/G1 ratios are compared to scrambled sequence OE.

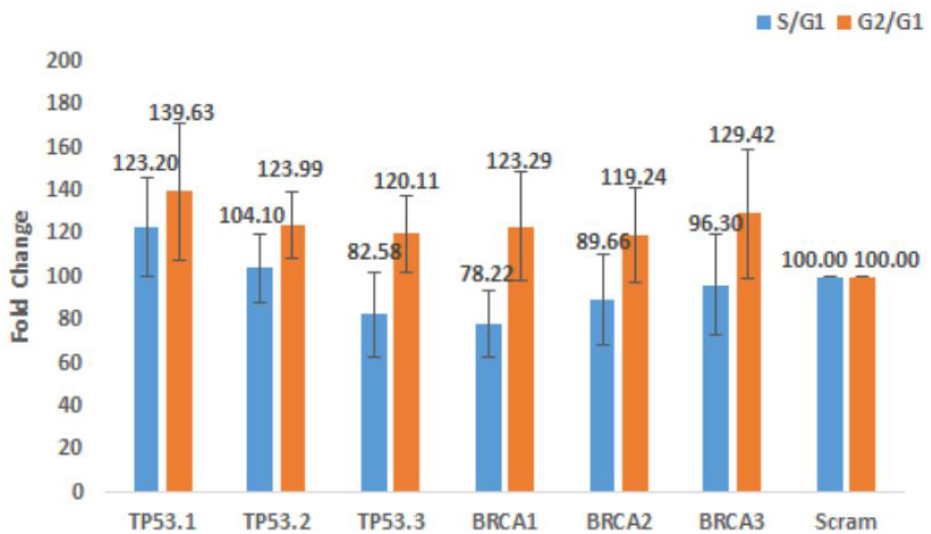

Cell-cycle analysis following 72h of RNA (OE) in the HeLa cell line.

S/G1 or G2/G1 ratios are compared to scrambled sequence OE. BrdU and 7-AAD expressions were detected by Gallios FACS machine and determined by Kaluza software version 1.3. Paired Student’s t-test was used for the statistical analysis (n=3).

Statistical analysis methods: Data are presented as mean ± SEM. p-values were calculated using a paired t-test, with p <0.05 considered significant (*<0.05, **<0.005).

## Future Development

Evidence supports education of NK cells using Codondex cell selections for autologous therapies. Papers supporting NK and p53 relationships follow;

In studies P53 is central:

**G**iven the role of Trp53 as a tumor suppressor gene and its associated mutations’ causative role in cancer, it is possible that patients harboring specific polymorphisms in Trp53 may also have alterations in NK cell functional maturation and subsequent defects in tumor immunosurveillance. Hence, investigation of the proportion of mNK cell subsets in patients harboring Trp53 polymorphisms would be of great interest. To that effect, various polymorphisms in both validated and putative p53 response elements, which would alter p53 binding and consequently the transcription of the given gene, have been identified in humans^17^.

**I**n this study, we found that p53 can sensitize tumor-killing susceptibility to GzmK or NK cell-mediated cytolysis. p53 associates with GzmK and is a physiological substrate of GzmK. p53 was processed to three cleavage products at Lys24 and Lys305. These three cleavage products harbor strong proapoptotic activities that amplify the proapoptotic action of p53 to potentiate tumor-killing sensitivity^18^.

Here we suggest NK surveillance is at least a response to changes in target homeostasis of p53 RNA isoforms, most likely via exosomal RNA concentration density^19 20^. NK cells deposit exosomes containing RNA, granzymes B and K, of which p53 is a physiological substrate and perforin. Exosomes fill the immunological synapse, bind receptors, release membrane penetrating perforin and promote lysis when granzymes cleave upregulated P53, BID, Caspase upregulate FAS and other pathways^*^^21 22^. This causes Bax and associated mitochondrial reactions, membrane permeabilization, pressure loss, release of Cytochrome C, collapse of electron transport chain and apoptosis^23^.

**F**indings suggest differential Basal and Squamous cell carcinoma risk with centromeric and telomeric activating KIR, and selective pressure from this NK based inter-individual antigen variability drives p53 alteration in skin tumors^25^.

Codondex computations identify highly specific intronic biomarkers that are used to classify and screen cells for therapeutic application. Codondex iScore™ is the measure by which transcript pkmers of same gene transcripts obtain a representative position in a vector to screen and select cells that are experimentally confirmed to function in a prescribed manner.

RNAScope and ISH^26^ is a compelling example of a method to probe a patient sample, isolate intron and the translated protein for sequencing. The RNAScope provides visual markers in protein to be extracted for Mass Spectrometry and sequencing - which is a preference over mRNA sequencing because it represents the final protein product from which a signature is calculated for use in Codondex iScore™.

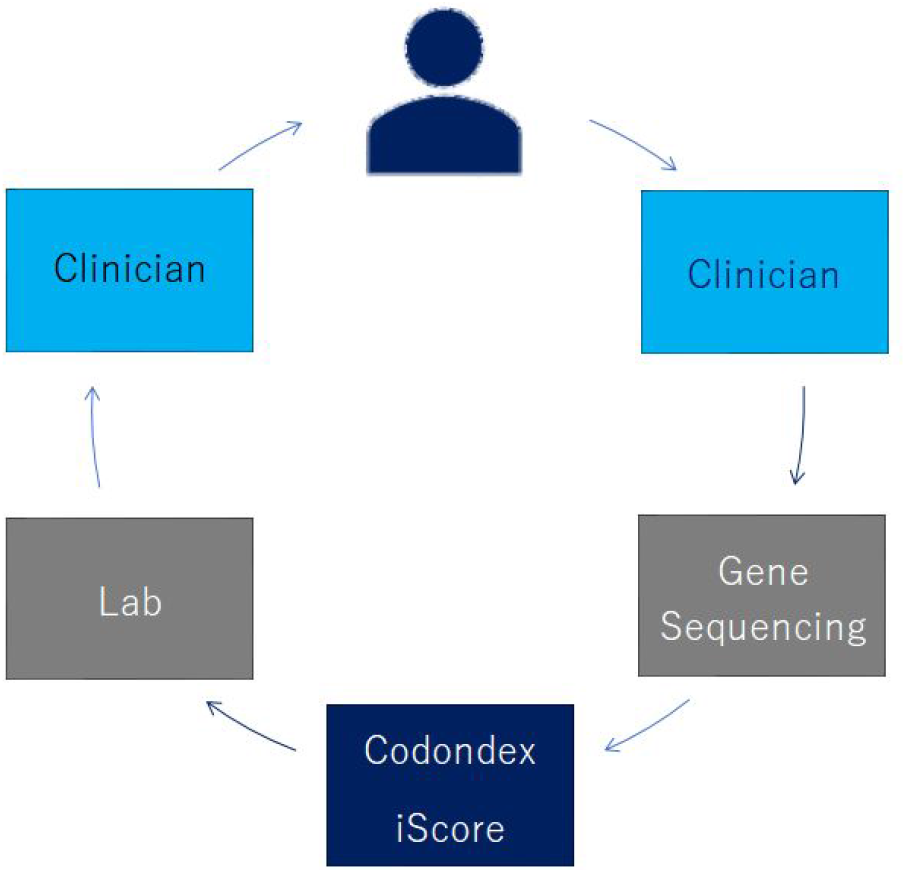

In the Codondex protocol multiple fragments of a tissue sample (or liquid biopsy) are separated^27^. Fresh cells are purified from each fragment and sequenced to determine a single fragment selection as follows;

1. RNASeq intron1 and sequence mRNA/protein
2. Compute Codondex iScore^™^ for each fragment
3. Select lead 28+ oligo target pkmers
  a. Identify fragment with highest leading pkmer count
4. Extract patient NK cells
  a. co-culture NK with selected fragment cells in whole blood/PMBC media
5. Extract NK from fragment
6. Introduce to patient

In a continuation;

1. NK are subjected to autologous expansion when cocultured with normal fibroblasts^28^
2. Immature and mature re-educated NK tested for cytotoxicity and immune response against target cells from pre-selected fragments ex vivo^29,30^.
3. NK dosage tested, applied in vivo

## About Codondex

Codondex and its parent company Precision Autology are the privately funded interests of Kevin Bermeister. Participating institutions in Sydney, Israel, London and Los Angeles are contributing to develop therapies combining Codondex i-Score computation using wholly autologous cell and blood selections from single patients.

## Footnotes

* (Restoring wild type p53 has been shown to restore effector NK function on tumor cells by induction of FAS^24^ and restoration of FAS apoptotic pathway.)

## References

1 Transposons, p53 and Genome Security Bhavana Tiwari, Amanda E. Jones, John M. Abrams

2 The Unusual Phylogenetic Distribution of Retrotransposons: A Hypothesis Jef D. Boeke, Department of Molecular Biology and Genetics, Johns Hopkins University School of Medicine, Baltimore, Maryland 21205, USA

3 Cytoplasmic intron sequence-retaining transcripts (CIRTs) can be dendritically targeted via ID element retrotransposons Peter T. Buckley, Miler T. Lee, Jai-Yoon Sul, Kevin Y. Miyashiro, Thomas J. Bell, Stephen A. Fisher, Junhyong Kim, and James Eberwine

4 High SINE RNA Expression Correlates with Post-Transcriptional Downregulation of BRCA1 Maureen Peterson, Vicki L. Chandler and Giovanni Bosco

5 Huergo-Zapico L, Acebes-Huerta A, López-Soto A, Villa-Álvarez M, Gonzalez-Rodriguez AP, Gonzalez S. Molecular Bases for the Regulation of NKG2D Ligands in Cancer. Frontiers in Immunology. 2014;5:106. doi:10.3389/fimmu.2014.00106.

6 Epstein-Barr virus-encoded microRNAs as regulators in host immune responses Man Wang, Fei Yu, Wei Wu, Yu Wang, Han Ding, Lili Qian Institute for Translational Medicine, Medical College of Qingdao University, Dengzhou Road 38, Qingdao 266021, China

7 A Novel Type of Non-coding RNA, nc886, Implicated in Tumor Sensing and Suppression Yong Sun Lee

8 p53 Upregulates Death Receptor 4 Expression through an Intronic p53 Binding Site Xiangguo Liu, Ping Yue, Fadlo R. Khuri and Shi-Yong Sun

9 A tissue-specific promoter derived from a SINE retrotransposon drives biallelic expression of PLAGL1 in human lymphocytes Claire E. L. Smith, Alexia Alexandraki, Sarah F. Cordery¤, Rekha Parmar, David T. Bonthron, Elizabeth M. A. Valleley*

10 Decoding ALS: from genes to mechanism J. Paul Taylor, Robert H. Brown Jr & Don W. Cleveland

11 Creation of the two isoforms of rodent NKG2D was driven by a B1 retrotransposon insertion C. Benjamin Lai, Ying Zhang, Sally L. Rogers and Dixie L. Mager*

12 Transposable elements 1 generate regulatory novelty in a tissue specific fashion Marco Trizzino, Aurélie Kapusta and Christopher D. Brown

13 Human NK Cells Are Alerted to Induction of p53 in Cancer Cells by Upregulation of the NKG2D Ligands ULBP1 and ULBP2 Sonja Textor, Nathalie Fiegler, Annette Arnold, Angel Porgador, Thomas G. Hofmann, and Adelheid Cerwenka

14 Direct Transcriptional Activation of Promyelocytic Leukemia Protein by IFN Regulatory Factor 3 Induces the p53 Dependent Growth Inhibition of Cancer Cells Tae-Kyung Kim, Joong-Seob Lee, Se-Yeong Oh, Xun Jin, Yun-Jaie Choi, Tae-Hoon Lee, Eun ho Lee, Young-Ki Choi, Seungkwon You, Yong Gu Chung, Jang-Bo Lee, Ronald A. DePinho, Lynda Chin, and Hyunggee Kim

15 Molecular mechanisms of natural killer cell activation in response to cellular stress CJ Chan, MJ Smyth and L Martinet

16 Each pkmer was ordered by frequency of their oligo repeats (>7) found for the transcript computation (iScore) and selected when for the next pkmer the most transcripts changed positions in a highly ordered vector (PiV).

17 An Unbiased Linkage Approach Reveals That the p53 Pathway Is Coupled to NK Cell Maturation Roxanne Collin, Charles St-Pierre, Lorie Guilbault, Victor Mullins-Dansereau, Antonia Policheni, Fanny Guimont-Desrochers, Adam-Nicolas Pelletier, Daniel H. Gray, Elliot Drobetsky, Claude Perreault, Erin E. Hillhouse and Sylvie Lesage

18 Ignition of p53 Bomb Sensitizes Tumor Cells to Granzyme K-Mediated Cytolysis Guoqiang Hua, Shuo Wang, Chao Zhong, Peng Xue and Zusen Fan

19 Extracellular vesicles released by CD40/IL-4 stimulated chronic lymphocytic leukemia cells confer altered functional properties to CD4+ T cells Dawn T. Smallwood, Benedetta Apollonio, Shaun Willimott, Larissa Lezina, Afaf Alharthi, Ashley R. Ambrose, Giulia De Rossi, Alan G. Ramsay and Simon D. Wagner

20 microRNAs responsive to A. actinomycetemcomitans and P. gingivalis LPS modulate expression of genes regulating innate immunity in human macrophages Afsar R. Naqvi, Jezrom B. Fordham, Asma Khan, and Salvador Nares

21 Ignition of p53 Bomb Sensitizes Tumor Cells to Granzyme K-Mediated Cytolysis Guoqiang Hua, Shuo Wang, Chao Zhong, Peng Xue and Zusen Fan

22 Targeted Cell-to-Cell Delivery of Protein Payloads via the Granzyme-Perforin Pathway Daniel J. Woodsworth, Lisa Dreolini, Libin Abraham, Robert A.

23 Granzyme B-Activated p53 Interacts with Bcl-2 To Promote Cytotoxic Lymphocyte-Mediated Apoptosis Thouraya Ben Safta, Linda Ziani, Loetitia Favre, Lucille Lamendour, Gwendoline Gros, Fathia Mami-Chouaib, Denis Martinvalet, Salem Chouaib and Jerome Thiery

24 Potentiation of a Tumor Cell Susceptibility to Autologous CTL Killing by Restoration of Wild-Type p53 Function Jérôme Thiery, Guillaume Dorothée, Hedi Haddada, Hamid Echchakir, Catherine Richon, Rodica Stancou, Isabelle Vergnon, Jean Benard, Fathia Mami-Chouaib and Salem Chouaib

25 Skin cancer risk is modified by KIR/HLA interactions that influence the activation of natural killer immune cells Karin A. Vineretsky, Margaret R. Karagas, Brock C. Christensen, Jacquelyn K. Kuriger-Laber, Ann E. Perry, Craig A. Storm and Heather H. Nelson

26 https://acdbio.com/science/applications/research-solutions/complementary-rna-protein-analysis https://www.youtube.com/watch?v=CPgZEf6EyJQ http://cnpg.comparenetworks.com/348498-Watch-Webinar-Diagnostic-Applications-of-the-RNAscope-RNA-ISH-Technology/

27 Expansion and Homing of Adoptively Transferred Human NK Cells in Immunodeficient Mice Varies with Product Preparation and In Vivo Cytokine Administration: Implications for Clinical Therapy Jeffrey S. Miller, Cliona M Rooney, Julie Curtsinger, Ron McElmurry, Valarie McCullar, Michael R. Verneris, Natalia Lapteva, David McKenna, John E. Wagner, Bruce R. Blazar, and Jakub Tolar

28 Survival, Retention, and Selective Proliferation of Lymphocytes Is Mediated by Gingival Fibroblasts Carolyn G. J. Moonen, Sven T. Alders, Hetty J. Bontkes, Ton Schoenmaker, Elena A. Nicu, Bruno G. Loos and Teun J. de Vries

29 NK cell education via nonclassical MHC and non-MHC ligands Yuke He and Zhigang Tian

30 Mature natural killer cells reset their responsiveness when exposed to an altered MHC environment Nathalie T. Joncker, Nataliya Shifrin, Frédéric Delebecque, and David H. Raulet

